# Arms race of physical defences: Hooked trichomes of *Macaranga* ant-plants kill lycaenid caterpillars, but one specialist has a counter-defence

**DOI:** 10.1101/2025.01.28.635112

**Authors:** Ritabrata Chowdhury, Ulmar Grafe, Faizah Metali, Walter Federle

## Abstract

The co-evolution of insects and chemical plant defences has been described as an arms-race, but it is unclear if physical plant defences can produce similar outcomes. Here we report a previously unknown interaction from the mutualism between ants and *Macaranga* trees. Although *Macaranga* trees are well protected against herbivory by aggressive ants, caterpillars of the genus *Arhopala* (Lepidoptera: Lycaenidae) can feed on the leaves by appeasing the ants with nectar-like secretions. One ant-plant species, *M. trachyphylla*, bears hooked trichomes on its green surfaces. When placed on *M. trachyphylla* stems or petioles, *Arhopala* caterpillars associated with other *Macaranga* species (*A. major, A. dajagaka* and *A. zylda*) were quickly arrested by the sharp trichomes which pierced their cuticle, resulting in rapid loss of blood and death. In striking contrast, *A. amphimuta* caterpillars, which occur naturally on *M. trachyphylla*, could easily walk over the hooked trichomes without any injury. As hooked trichomes are a novel trait within *Macaranga*, this interaction provides an example of *de novo* evolution of a physical plant defence, which in turn has been overcome by a specialist herbivore. Our study suggests that physical plant defences can lead to evolutionary arms races similar to those for chemical defences.

## Introduction

Plants and insects have coexisted for more than 400 million years and over this period, they have developed complex interactions which play a critical role in terrestrial ecosystems (Bronstein et al. 2006; Endara et al. 2017; War et al. 2018). While some of these relationships are mutually beneficial, such as pollination or seed dispersal, the majority involve insects feeding of plants, and plants defending themselves against herbivory (War et al. 2018). The mutual adaptation of chemical plant defences and insect herbivores has been described as an evolutionary arms-race that has led to the diversification of both insects and plants (Ehrlich & Raven 1964; Endara et al. 2017). Do such arms-races also occur for physical defences?

One of the physical defences of plants against herbivores are trichomes, hair-like appendages produced by the epidermis. Trichomes can protect against herbivory by limiting insect access to leaf or stem tissues (Fordyce & Agrawal 2001; Kariyat et al. 2019; Kariyat et al. 2017), exposing insects to toxic or sticky substances (Hanley et al. 2007; Wagner 1991) or puncturing the insect’s cuticle (Gilbert 1971; Pillemer & Tingey 1976). Trichomes can also decrease the rate of tissue ingestion and thereby the herbivore’s growth (Kariyat et al. 2019; Kariyat et al. 2017), and prevent insect oviposition or reduce the adhesion of eggs to the plant surface (Haddad & Hicks 2000; Handley et al. 2005). Sharp hooked trichomes are considered one of the most effective forms of physical defence in plants. In *Passiflora* vines, they can injure or kill caterpillars by piercing their cuticle (Cardoso 2008; Gilbert 1971). Similarly, hooked trichomes on the leaves of common bean (*Phaseolus vulgaris*) can injure insects in different parts of their body, leading to their death by haemolymph loss or starvation (Pillemer & Tingey 1976; Szyndler et al. 2013; Xing et al. 2017).

Some insect herbivores have in turn evolved adaptations to deal with trichomes. These adaptations can be morphological, such as elongated legs to avoid contact with the sticky heads of glandular trichomes (Voigt et al. 2007) or tarsal hooks for anchoring to trichomes (Medeiros & Boligon 2007). Insects can also adapt behaviourally by walking “on tiptoe” between trichomes (Voigt et al. 2007), removing dense trichome covers before feeding on leaves (Shelomi et al. 2010), or spinning silk mats to move on trichome-bearing surfaces without being punctured (Rathcke & Poole 1975).

Here we report an interaction involving trichomes as a physical defence in ant-associated trees of the paleotropical genus *Macaranga* (Euphorbiaceae). Most *Macaranga* species are myrmecophilic and protect themselves against herbivory by attracting non-specific foraging ants, but ca. 30 species in South-East Asia are myrmecophytic and permanently house specialised ant partners nesting in their hollow stems. Although the ants efficiently keep away most herbivores, several species of *Arhopala* caterpillars (Lepidoptera: Lycaenidae) have outwitted the ants’ defences by providing them with nectar-like secretions from their myrmecophilous organs, and feed on *Macaranga* leaves (Maschwitz et al. 1984; Shimizu-Kaya et al. 2015). One *Macaranga* ant-plant species in Borneo, *M. trachyphylla*, has stems and petioles densely covered by hooked trichomes, giving the surfaces a sandpaper-like feel. To our knowledge, *M. trachyphylla* is the only one of more than 300 described *Macaranga* species for which this trait is known (Whitmore 1975). The functional significance of these trichomes is still unknown. We therefore investigated the effect of *M. trachyphylla* trichomes on *Arhopala* caterpillars.

## Results

*A. major* caterpillars attempting to walk on petioles or stems of *M. trachyphylla* were rapidly arrested by the sharp hooked trichomes (Fig.1A,B). As soon as they became stuck on a trichome, the caterpillars could only move about 1-2mm further. The sharp trichome tips pierced the caterpillars’ prolegs and/or abdomen and thorax (Fig.1C,D; Fig.2B,C; Supplementary Videos S1,S2), leading to the oozing out and rapid loss of haemolymph. All *A. major* caterpillars of the 1^st^ to 4^th^ instar (n=20) died very quickly, with death usually occurring within half an hour of the injury (Fig.3). The only *A. major* caterpillar not immediately killed on *M. trachyphylla* stems was from the final (5^th^) instar (1 out of 5 caterpillars; Fig.3); it was also arrested by the trichomes, but as the piercing occurred only in the prolegs, it did not die as a result of the interaction.

**Fig 1:**
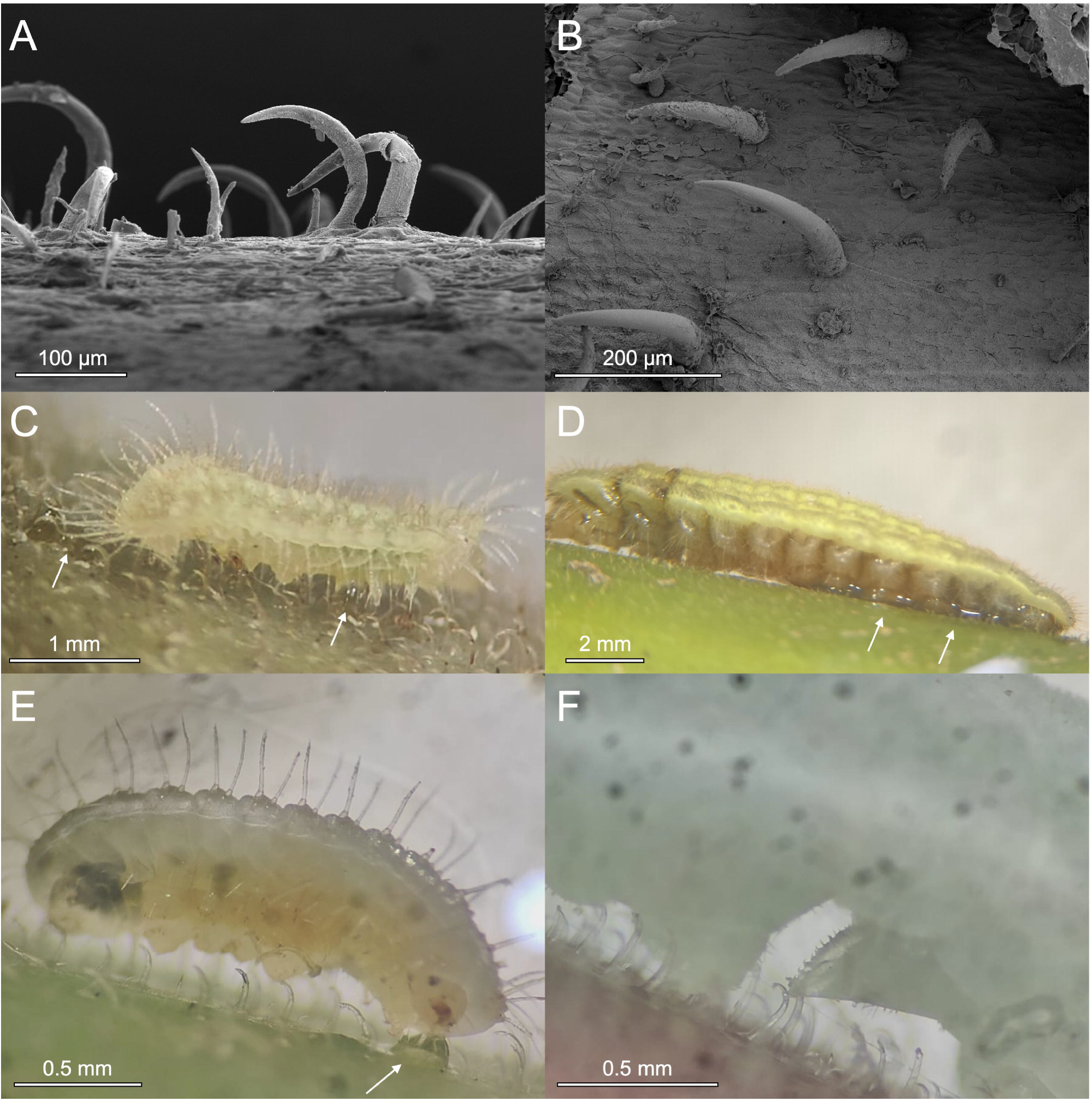
A,B) Scanning electron micrographs of hooked trichomes on the petiole (A) and stem (B) of *M. trachyphylla*; C) *Arhopala* major caterpillar (2nd instar) injured by *M. trachyphylla* trichomes, with haemolymph oozing from the wounds (arrows), D) same as C, but 4th instar; E) 1st-2nd instar A. dajagaka caterpillar, with haemolymph oozing from the wounds (arrow); and F) 3^rd^ instar A. zylda caterpillar, showing proleg footpad pierced by a hooked trichome.

**Fig 2:**
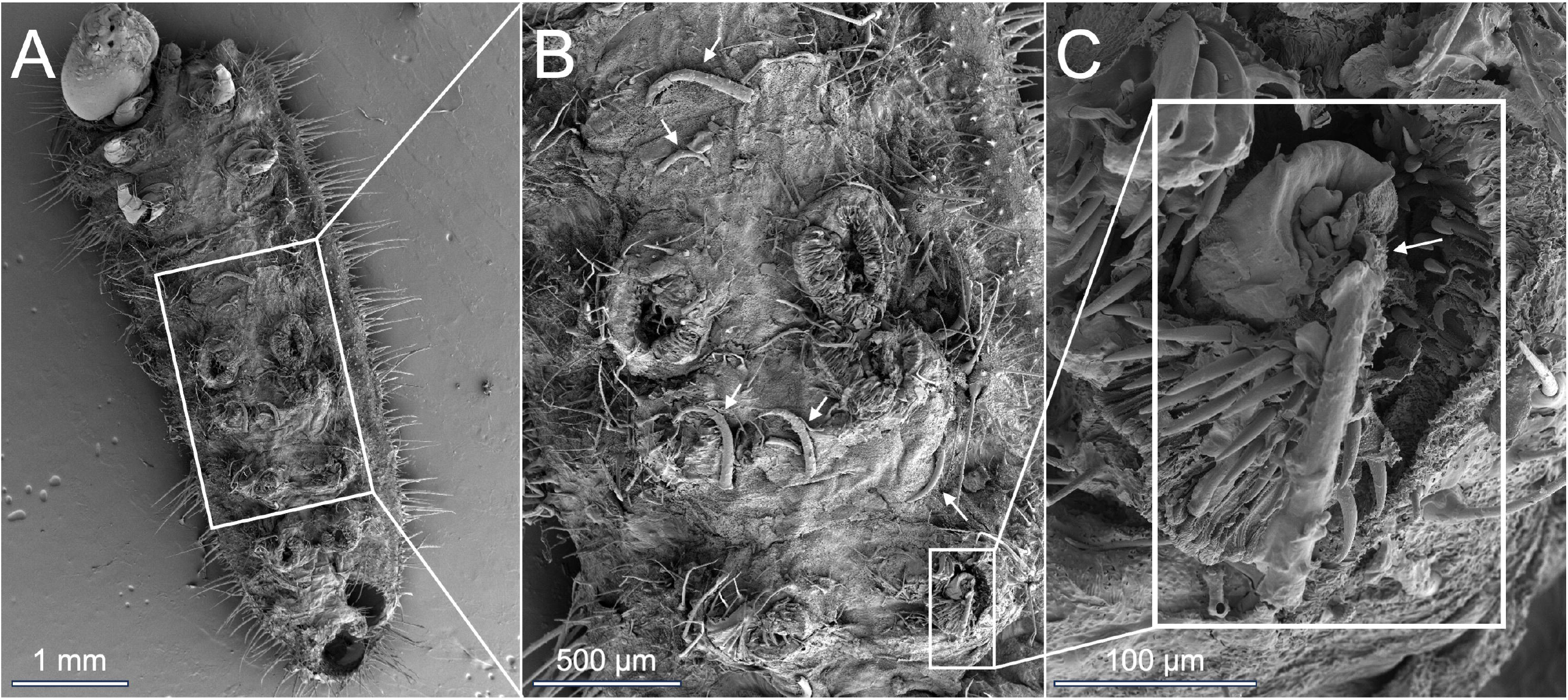
A-C) Scanning electron micrographs of ventral side of a 3^rd^-4^th^ instar *A. major* caterpillar after walking on the trichome-bearing stem surface of *M. trachyphylla*; B) & C) magnified images showing several broken trichomes (arrows in B) on the surface of the cuticle, and a hooked trichome piercing the cuticle of the 3^rd^ left proleg footpad (arrow in C).

**Fig 3:**
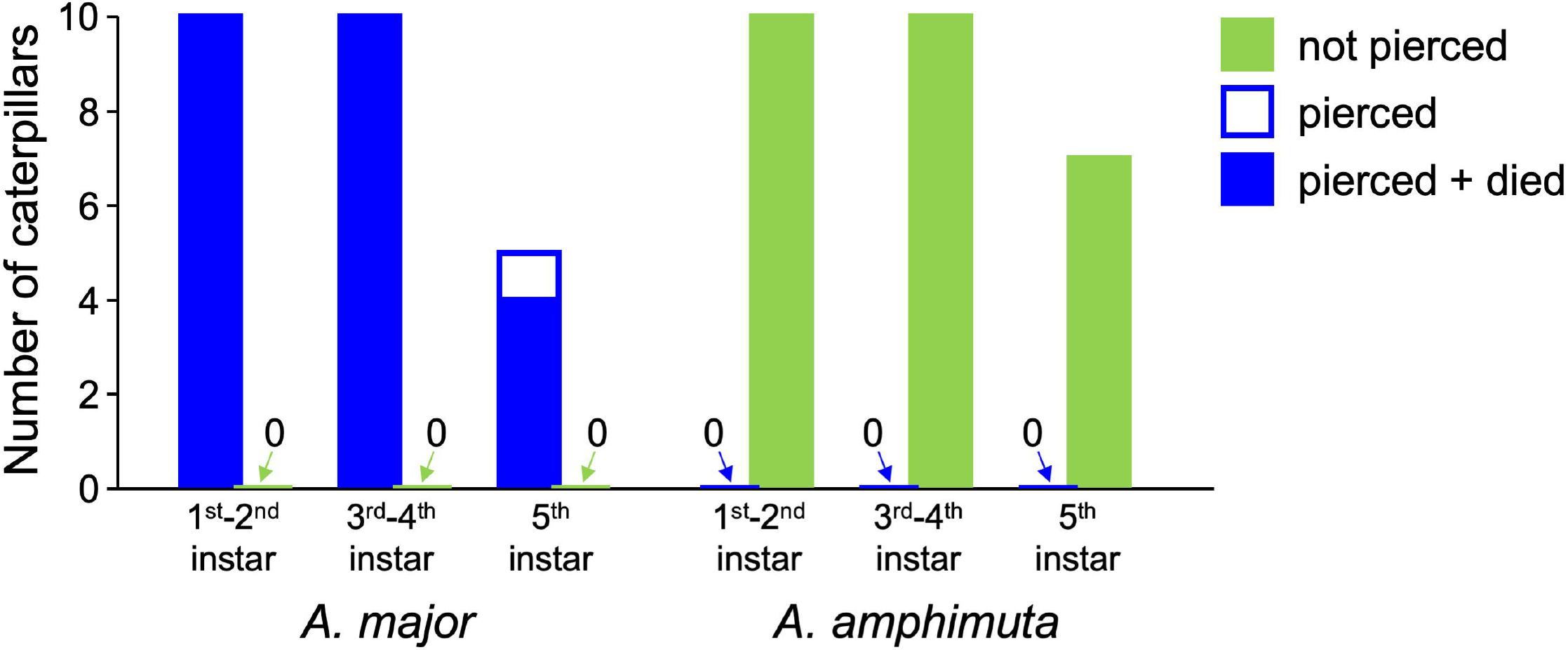
Effect of hooked trichomes on petioles and stems of *M. trachyphylla* on *A. major* and *A. amphimuta* caterpillars.

We found very similar effects in *A. zylda* and *A. dajagaka* caterpillars, although fewer specimens were available from these species (Fig.1E,F). All the caterpillars were arrested by the trichomes; *A. dajagaka* died due to loss of haemolymph, whereas the *A. zylda* caterpillars (Supplementary Video S3) could not move more than 1-2 mm, but did not die as they were only injured in their prolegs.

In striking contrast, *A. amphimuta* caterpillars of all instars were able to move easily over the hooked trichomes on *M. trachyphylla* stems and petioles, and no injuries were observed in any of the caterpillars (Fig.3; Supplementary Video S4). We observed that the prolegs or the body wall often caught and pulled on the hooked trichomes, but the trichomes did not puncture the cuticle. The caterpillars sometimes raised their prolegs high to free them from the trichomes, but we also observed this behaviour in the other *Arhopala* species tested. *A. amphimuta* caterpillars were always able to free their prolegs from the trichomes without being pierced. In contrast to *M. trachyphylla* stems and petioles, the leaves of *M. trachyphylla* were harmless for all the *Arhopala* species, probably because they contained a lower density of hooked trichomes; all caterpillars could walk on them without being injured.

We tested whether *A. major* caterpillars would be able to cope with the mutualistic *Crematogaster* plant-ants living on *M. trachyphylla* trees. When placed on the youngest leaves, the caterpillars were mostly ignored or attended by the ants (52% and 34% of caterpillar-ant interactions, respectively); they were only occasionally attacked (14% of interactions). However, when placed on the stem, all *A. major* caterpillars (n=10) became stuck in the trichomes within only 5 minutes, were punctured by them and started to lose haemolymph. Once the caterpillars were injured, the ants’ behaviour changed; there were quickly many ants that attacked and removed the caterpillar (n=10), probably due to the injury and because it was no longer able to use its myrmecophilous organs.

## Discussion

Our results show that the hooked trichomes of *M. trachyphylla* act as a powerful physical defence against *A. major, A. zylda* and *A. dajagaka* caterpillars, and that the *Arhopala* species naturally occurring on *M. trachyphylla, A. amphimuta*, is resistant to them. The sharp trichomes pierce the caterpillars’ cuticle, leading to rapid blood loss and death mostly occurring within minutes.

The action of the hooked trichomes of *M. trachyphylla* against *Arhopala* caterpillars and the presence of a species that is immune to this physical defence represent a striking evolutionary convergence with the interaction of *Passiflora* trichomes and *Heliconius* caterpillars discovered by Gilbert (1971). *Heliconius* caterpillars are also stopped, punctured and killed by the sharp hooked trichomes on *Passiflora* leaves, and there is one species, *H. charithonia*, that is resistant and capable of feeding on trichome-bearing plants (Cardoso 2008; MacDougal 1994). However, the *Arhopala-M. trachyphylla* interaction discovered here differs in some important ways from the *Passiflora-Heliconius* system:

Firstly, *M. trachyphylla* trees are ant-plants permanently inhabited by colonies of *Crematogaster* ants. These ants provide a powerful biotic defence that the herbivorous *Arhopala* caterpillars have to cope with. The five closely related *Arhopala* species associated with *Macaranga* (which form the amphimuta amphimuta subgroup, Megens et al. 2005), each occur on a non-overlapping set of *Macaranga* species. It is still unclear whether this host specificity is based on ant defences, or on chemical or physical plant traits. We found that *A. major* caterpillars were not strongly rejected by the plant-ants living on *M. trachyphylla*, and similar observations have been reported for *A. zylda* and *A. dajagaka* (Inui et al. 2015). It is also unlikely that the specificity of *A. major* for *M. gigantea* and absence on *M. trachyphylla* is due to defensive plant chemicals, because *A. major* caterpillars can feed on *M. trachyphylla* leaves and successfully pupate (Shimizu-Kaya et al. 2013). Our findings show that the absence of *A. major, A. zylda* and *A. dajagaka* caterpillars on *M. trachyphylla* can be fully explained by the sharp, hooked trichomes on the stems and petioles.

A second difference to *Passiflora* is that the trichome defence of *M. trachyphylla* is restricted to the petioles and stem. *A. major* caterpillars can feed on *M. trachyphylla* leaves in the laboratory (Shimizu-Kaya et al. 2013), but under natural conditions, the trichomes on petioles and stem are sufficient to prevent them from completing their development. Assuming an *A. major* egg is laid on a young leaf of *M. trachyphylla*, the emerging young caterpillar will at some point have to move to the next leaf via the petiole and stem. In this process, the caterpillar will be trapped and killed by the trichomes.

It has long been proposed that there is a co-evolutionary arms race between the chemical defences of plants and insect counter-adaptations (Ehrlich & Raven 1964; Endara et al. 2017; Howe & Jander 2008). Until now, it is unclear whether physical defences can trigger similar arms-races. The interaction of hooked trichomes in *M. trachyphylla* and *Arhopala* caterpillars might represent one of the first examples of such an arms-race. As the main anti-herbivore defence of *Macaranga* trees via their ant partners has already been overcome by *Arhopala* caterpillars (Inui et al. 2015; Maschwitz et al. 1984), the isolated occurrence of hooked trichomes in *M. trachyphylla* suggests that they evolved as a new defence to cope with heavy *Arhopala* herbivory. In turn, the ability of *A. amphimuta* caterpillars to walk over the hooked trichomes of *M. trachyphylla* without sustaining any injuries could represent an adaptation in response to this new defence. However, this arms-race scenario is still speculative, as the detailed mechanisms underlying the resistance of *A. amphimuta* are still unknown, and it is equally possible that the relevant traits evolved as pre-adaptations for some other functions.

*A. amphimuta* caterpillars occur not only on *M. trachyphylla* but also on several other *Macaranga* host plants that are free of hooked trichomes. Since *M. trachyphylla* is endemic to Borneo, populations of *A. amphimuta* living on M. bancana and M. hullettii on the Malay peninsula have never encountered hooked trichomes. Future research on these populations could shed light on whether the resistance of *A. amphimuta* to hooked trichomes is indeed a counter-adaptation or a pre-adaptation that has allowed this species to survive on *M. trachyphylla*.

## Methods

### Field site

*Arhopala* caterpillars associated with *Macaranga* trees were studied in the Andulau Forest Reserve and the Ulu Temburong National Park in Brunei from May to June 2023 and from July to September 2024. *A. major* caterpillars were collected from M. gigantea trees (n=25), *A. amphimuta* caterpillars from M. bancana (n=17), *M. trachyphylla* (n=5), M. hullettii (n=4) and M. aëtheadenia (n=1; to our knowledge, this is the first report of M. aëtheadenia as host plant of A. amphimuta). A. dajagaka was collected from M. rufescens (n=1), and A. zylda from M. beccariana (n=3).

### Trichome morphology

The morphology of hooked trichomes from leaves, stems and petioles of *M. trachyphylla* was studied under a Leica EZ4 stereomicroscope (Leica Microsystems, Wetzlar, Germany) and using scanning electron microscopy. For electron microscopy, leaf, petiole and stem samples were first fixed overnight in formalin-alcoholic solution (4% v/v formaldehyde, 70% v/v ethanol), followed by gradual dehydration in an ethanol series (70%, 90% and absolute ethanol). The samples were then dried in a Quorum E3100 critical point dryer, sputter-coated with 20 nm of Platinum (Quorum K575X, Quorum Technologies Ltd., East Grinstead, UK) before viewing them under the scanning electron microscope (Tescan MIRA3 FEG; TESCAN UK Ltd., Cambridge, UK) at 5eV.

### Effect of hooked trichomes on *Arhopala* caterpillars

Observations of caterpillars placed on leaves, petioles and stems of *M. trachyphylla* were made in Brunei under a Leica EZ4 stereomicroscope (Leica Microsystems, Wetzlar, Germany) and recorded with the camera of a OnePlus 11R mobile phone (OnePlus Technology Co., Ltd, China) attached to the microscope by an adapter (Celestron 81035). 25 *A. major* caterpillars were studied (n=10 1^st^-2^nd^ instar, n=10 3^rd^-4^th^ instar and n=5 5^th^ instar and prepupa), as well as 27 *A. amphimuta* caterpillars (n=10 1^st^-2^nd^ instar, n=10 3^rd^-4^th^ instar and n=7 5^th^ instar and prepupa), three instar A. zylda (n=1 3^rd^, n=1 4^th^ instar and n=1 5^th^ instar) and one 1^st^-2^nd^ instar A. dajagaka. Each caterpillar was carefully placed on the surface using soft tweezers and then observed and video-recorded for 10 mins under the stereo microscope. One 3^rd^-4^th^ instar caterpillar of *A. major* was freeze-dried after interacting with hooked trichomes on *M. trachyphylla* and viewed under the scanning electron microscope (see above) to visualise details of injuries.

### Ant behaviour towards *Arhopala* caterpillars

Previous work showed that *A. major* caterpillars are able to feed on *M. trachyphylla* leaves under ant-free conditions in the laboratory, where they reached pupation at a rate even higher than on leaves of their natural host M. gigantea (Shimizu-Kaya et al. 2013). We therefore tested whether *A. major* caterpillars are accepted by the *Crematogaster* plant-ants living on *M. trachyphylla*. The ants’ behavioural responses were studied by placing *A. major* caterpillars on ant-inhabited *M. trachyphylla* plants (as in Inui et al. 2015). 2^nd^, 3^rd^ or 4^th^ instar *A. major* caterpillars were collected from M. gigantea in the field and kept on young ant-free leaves of their host plant in a plastic container in the laboratory for up to 2 days before the introduction. For the experiments, young *M. trachyphylla* trees of 1.5–2.0 m height with low herbivory damage were chosen and one caterpillar was placed on the abaxial surface of the youngest or second-youngest visible leaf of the tree. The ants’ responses to the introduced larva were recorded for 10 min after the larva was approached for the first time by a worker of the resident ant colony. The reactions of ants that approached the caterpillar were divided into the following three categories (only one interaction was scored per ant visiting the leaf): 1) ignoring: no ants attacked or touched the caterpillar; 2) attending: ants touched the caterpillar without biting near the myrmecophilous glands on the posterior-dorsal side and sometimes harvested nectar; 3) attacking: ants bit the caterpillar. Each caterpillar and each ant colony (tree) were only used once. At the end of the trials, the caterpillar, if not already killed by the ants, was placed on the first internode of the stem and the caterpillar’s movements and ant behaviour were observed qualitatively for a further 10 minutes.

## Acknowledgements

We would like to thank Mr Federick Valentino bin Yahau for his assistance during this study and Dr Heather Greer, Department of Chemistry, University of Cambridge, for her help with Scanning Electron Microscopy. RC was funded by R. O. Whyte PhD Studentship and the Trinity Henry Barlow Studentship from the University of Cambridge, UK, the Varley-Gradwell Travelling Fellowship in Insect Ecology from the University of Oxford, UK, the Zoology Fieldwork Fund from the University of Cambridge, UK, and the Professor Hering Memorial Research Fund from the British Entomological and Natural History Society, UK. Furthermore, we would like to thank the Faculty of Science, and the Kuala Belalong Field Studies Centre, Universiti Brunei Darussalam for logistical support, the Forestry Department in Brunei for issuing the forest entry permits, collection permits and export permits and the Department of Agriculture and Agrifood, Brunei for issuing the phytosanitary certificate.

## Author contributions

Conceptualization: RC, WF; Methodology, Investigation and Visualization: RC; Supervision: WF, UG, FM; Writing – original draft: RC; Writing – review & editing: RC, WF, FM, UG Competing interests: Authors declare that they have no competing interests

## Supplementary Video legends

**Supplementary Video S1:** 2^nd^ instar *Arhopala* major caterpillar being injured by *M. trachyphylla* trichomes, with haemolymph oozing from the wound

**Supplementary Video S2:** 3^rd^ instar *A. major* caterpillar being injured by *M. trachyphylla* trichomes, with haemolymph oozing from the wound

**Supplementary Video S3:** 3^rd^ instar *A. zylda* caterpillar, showing proleg footpad being pierced by a hooked trichome

**Supplementary Video S4:** 3^rd^ instar *A. amphimuta* caterpillar walking over hooked trichomes on *M. trachyphylla* stem without being injured

